# The sodium-glucose cotransporter 2 inhibitor tofogliflozin induces vasodilation by activating Kv channels, the SERCA pump, and the sGC/cGMP pathway

**DOI:** 10.1101/2024.12.09.627651

**Authors:** Wenwen Zhuang, Minju Park, Junsu Jeong, Hye Ryung Kim, YeEun Jang, Mi Seon Seo, Jin Ryeol An, Hongzoo Park, Won-Kyo Jung, Il-Whan Choi, Won Sun Park

## Abstract

**OBJECTIVE:** Tofogliflozin is a sodium-glucose cotransporter 2 (SGLT2) inhibitor widely used to treat T2DM, but it also exhibits cardio-protective effects. This study investigated the vasodilatory action of tofogliflozin using rabbit femoral artery rings pre-contracted with phenylephrine.

**APPROACH AND RESULTS:** The femoral artery quickly separated from the rabbit and fix it to the organ bath chamber. Subsequently, administer an inhibitor that modulates vascular tension in the rings or remove the endothelium to assess its impact on vasodilation. The results showed the concentration-dependent induction of vasodilation by tofogliflozin, a response that remained unchanged following endothelial removal, pretreatment with the nitric oxide synthase (NOS) inhibitor L-NAME, or the inhibition of low- and intermediate-conductance Ca^2+^-activated K^+^ channels (SK_Ca_ and IK_Ca_) using apamin in combination with TRAM-34. Furthermore, pretreatment with the voltage-dependent K^+^ (Kv) channel inhibitor 4-AP reduced the vasodilatory effects of tofogliflozin whereas pretreatment with the ATP-sensitive K^+^ (K_ATP_) channel inhibitor glibenclamide or the large-conductance Ca^2+^-activated K^+^ (BK_Ca_) channel inhibitor paxilline did not. Notably, our findings indicated that Kv7.X, rather than Kv1.5 or Kv2.1, is the primary Kv subtype involved in tofogliflozin-induced vasodilation. The vasodilatory effects of tofogliflozin were also significantly inhibited in femoral arterial rings pretreated with the sarco/endoplasmic reticulum Ca^2+^-ATPase (SERCA) pump inhibitors thapsigargin and cyclopiazonic acid (CPA). Tofogliflozin-induced vasodilation was unaltered in arterial rings exposed to the adenylyl cyclase inhibitor SQ 22536, the protein kinase A (PKA) inhibitor KT 5720, and the protein kinase G (PKG) inhibitor KT 582 whereas it was effectively reduced by the soluble guanylyl cyclase (sGC) inhibitor ODQ.

**CONCLUSIONS:** These findings suggest that tofogliflozin-induced vasodilation is mediated by the activation of the SERCA pump, the sGC/cGMP pathway, and Kv channels, but not the PKA signaling pathway, other K^+^ channels, or endothelium-dependent mechanisms.

## INTRODUCTION

Diabetes is a major global health challenge, with type 2 diabetes mellitus (T2DM) comprising ∼90% of all cases^1^. Numerous clinical studies have demonstrated a high prevalence of cardiovascular issues among T2DM patients, including hypertension, obesity, and dyslipidemia^2^. Therefore, the management of hyperglycemia in T2DM patients should include not only the control blood glucose levels but also the reduction of cardiovascular risk, which has driven the development of antidiabetic therapies with cardioprotective properties^3^.

Sodium-glucose cotransporter 2 (SGLT2) inhibitors represent a notable advancement in T2DM treatment, with significant implications for the management of cardiovascular disease^4^. By promoting glucose excretion via renal filtration, SGLT2 inhibitors enhance glycemic control, thereby facilitating weight loss and reducing the combined risks of cardiovascular mortality and renal events^5–7^. The SGLT2 inhibitor tofogliflozin was initially approved in Japan in 2014 for the treatment of adults with T2DM, based on its ability to improve glycemic control and its beneficial cardiovascular and metabolic effects^8, 9^. A subsequent study showed that T2DM patients treated with tofogliflozin had improved serum lipid profiles and reduced body weight^10^. Additional benefits of tofogliflozin include cardiovascular protective effects, particularly against hypertension and heart failure^11^. However, the precise mechanisms underlying tofogliflozin-mediated cardiovascular protection remain poorly understood.

Vascular tension is regulated by the interaction of ion channels and intracellular signaling pathways. Ion channels, especially potassium (K^+^) channels, control membrane potential and are thus key regulators of vascular tone^12^. Four types of K^+^ channels are expressed in most vascular smooth muscle cells (VSMCs). Among these, voltage-dependent K^+^ (Kv) channels regulate steady-state membrane potential and vascular tension. Their activation leads to membrane hyperpolarization, which in turn relaxes vascular smooth muscle by reducing Ca^2+^ influx^13, 14^. This negative feedback mechanism is also regulated by intracellular signaling pathways, both directly and indirectly. For example, the sarco/endoplasmic reticulum Ca^2+^-ATPase (SERCA) pump indirectly maintains membrane hyperpolarization by lowering cytoplasmic Ca^2+^ levels, supporting Kv channel activation and relaxation^15^. The soluble guanylyl cyclase (sGC)/cyclic guanosine monophosphate (cGMP) pathway directly enhances Kv channel activity through protein kinase G (PKG), further stabilizing hyperpolarization^16^. Alterations in the function of these signaling complexes are closely associated with diseases affecting the cardiovascular system, including diabetes, hypertension, and atherosclerosis^17^. Given the physiological importance of Kv channels in vascular function, their contribution and that of related signaling pathways to the cardiovascular protective effect of tofogliflozin merit further investigation.

This study investigated the mechanisms underlying the vasodilatory effects of tofogliflozin and demonstrated the involvement of SERCA pumps, sGC/cGMP pathways, and Kv channels. Our findings provide a new perspective for interpreting the clinical effects of tofogliflozin and suggest that it should be investigated for potential clinical applications beyond diabetes.

## MATERIALS AND METHODS

### Animals and femoral artery tension measurement

New Zealand white rabbits (8 weeks old, male, body weight 2.0–2.5 kg) were purchased from the Daehanbiolink Co. (Eumseong-gun, South Korea) and maintained at a constant temperature (25 ± 1°C), 50% humidity, and a standard 12-h light/12-h dark cycle. Standard chow and water were provided *ad libitum*. All animal experiments were performed in accordance with National Institutional of Health guidelines and approved by the Institutional Animal Care and Use Committee of Kangwon National University (approval no. KW-210512-2). The rabbits were anesthetized with heparin (100 U/kg) and sodium pentobarbitone (40 mg/kg) and allowed to rest for 3–5 min. The femoral artery was then quickly isolated and transferred to cold normal Tyrode’s solution (NT). Adipose tissue and residual blood in the femoral artery were promptly removed, and the artery was cut into ∼10-mm lengths. Two L-shaped stainless-steel wires were inserted into each arterial ring and connected with a force–displacement transducer (FORT25; WPI, Sarasota, FL, USA). The rings were then incubated in an organ bath filled with an oxygenated (95% O_2_ and 5% CO_2_) physiological salt solution (PSS) and maintained at a resting tension of 1.8–2.0 g for over 1 h at 37°C. Changes in isometric tension were measured using a force–displacement transducer. The signal was digitalized at 1 kHz using the Power Lab 4/35 data acquisition system (AD Instruments, Colorado Springs, CO, USA) and recorded with Lab Chart Pro v8.0 software. In experiments using endothelium-denuded femoral rings to assess endothelial dependency, the endothelium was removed by injecting air bubbles intraluminally over 40 times. Successful removal was confirmed using acetylcholine (1 μM). Arterial viability was tested by applying a high-K^+^ (80 mM) solution prior to starting the experiments.

### Blood pressure measurement

Blood pressure was measured using a noninvasive monitoring system (Bionics Co., Ltd., Seoul, South Korea) and 30-mm-wide cuffs without anesthesia. The rabbits were stabilized for 30 min with the cuff wrapped around the brachial artery. Blood pressure levels were assessed 2 h after the injection of tofogliflozin (1 mg/kg, corresponding to a human dose of 20 mg) into the ear vein, to allow measurement of the maximum plasma concentration.

### Reagents

The NT solution used for vessel preparation was composed of 5.4 mM KCl, 143 mM NaCl, 0.7 mM MgCl_2_, 1.6 mM CaCl_2_, 5 mM HEPES, 0.66 mM NaH_2_PO_4_, and 13 mM glucose; the pH was adjusted to 7.4 with NaOH. The PSS used for tension measurements was composed of 4.8 mM KCl, 144 mM NaCl, 25.6 mM NaHCO_3_, 1.2 mM KH_2_PO_4_, 1.8 mM CaCl_2_, 1.5 mM MgSO_4_, and 13 mM glucose; the pH was adjusted to 7.4 with NaOH. Both 4-aminopyridine (4-AP) and phenylephrine (Phe) were purchased from Sigma Chemical Co. (St. Louis, MO, USA). Tofogliflozin, glibenclamide, paxilline, SQ 22536, KT 5720, ODQ, KT5823, cyclopiazonic acid (CPA), thapsigargin, L-NAME, apamin, DPO-1, stromatoxin-1, linopirdine, and acetylcholine were purchased from Tocris Cookson (Ellisville, MO, USA). 4-AP, Phe, apamin, and stromatoxin-1 were dissolved in distilled water. Glibenclamide, paxilline, SQ 22536, KT 5720, ODQ, KT 5823, CPA, thapsigargin, L- NAME, DPO-1, linopirdine, and acetylcholine were dissolved in dimethyl sulfoxide (DMSO). The final concentration of DMSO was < 0.1%; DMSO at this concentration had no significant effect on Phe/drug-induced vasoconstriction or vasodilation.

### Data analysis and statistics

All data were analyzed using OriginPro-8 software (Microcal Software, Inc., Northampton, MA, USA). Statistical analyses were performed using GraphPad Prism v8.0 (GraphPad Software, La Jolla, CA, USA). The results are presented as the means ± standard error of the mean (SEM). Mann–Whitney *U* tests were used to evaluate statistical significance, at a threshold of *P* < 0.05.

## RESULTS

### Vasodilatory effects of tofogliflozin on Phe-induced pre-contracted femoral arterial rings

The effects of tofogliflozin on the rabbit femoral artery were investigated by pre-contracting femoral arterial rings with 1 μM Phe and then exposing them to increasing concentrations of tofogliflozin (1, 3, 10, 30, 50, 100, 300, and 500 μM). The vasodilatory effect of tofogliflozin on the femoral artery increased significantly at higher drug concentrations, in a concentration-dependent manner (Figure 1A). For example, treatment with 50 and 100 μM tofogliflozin increased vasodilation by 46.88% and 84.33%, respectively (Figure 1B).

**Figure 1.**
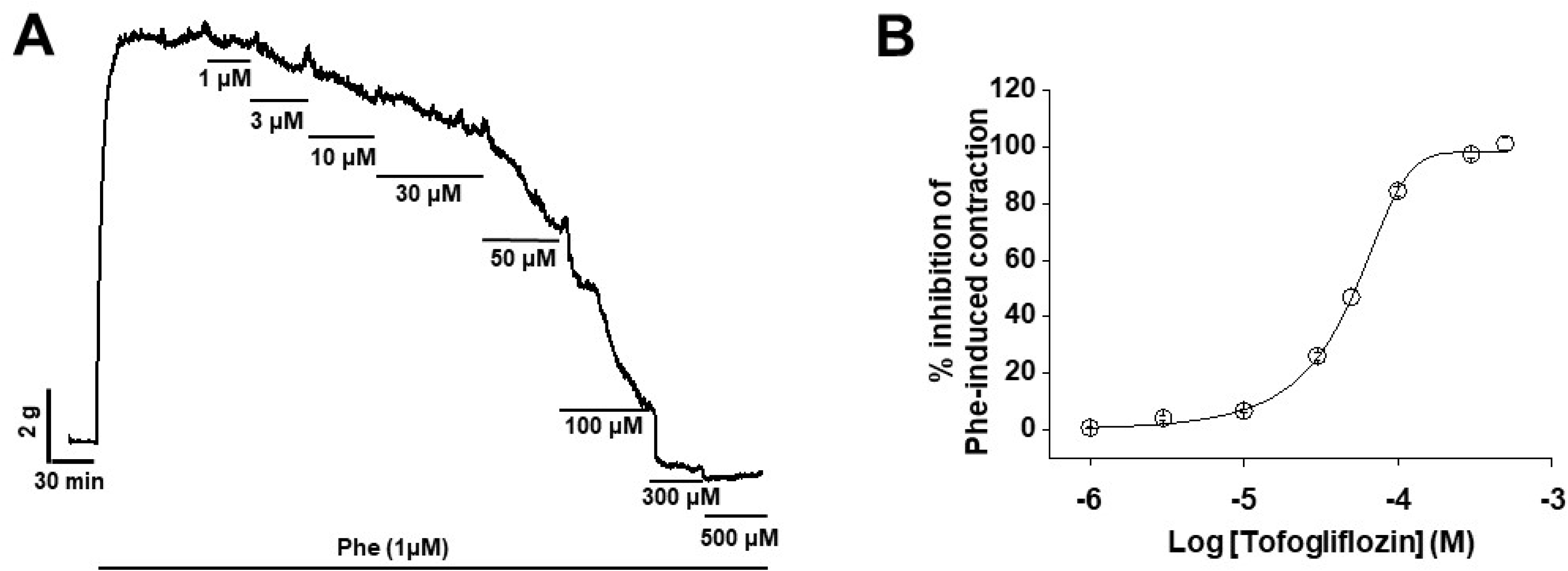
Effect of tofogliflozin on rabbit femoral arterial rings. (A) Vasodilatory effects of tofogliflozin at concentrations of 1, 3, 10, 30, 50, 100, 300, and 500 μM on femoral arterial rings pre-contracted with phenylephrine. (B) Concentration-dependent curve of tofogliflozin-induced vasodilation (*n* = 9; all n values refer to femoral arterial rings isolated from different rabbits).

### The role of K^+^ channel inhibitors in tofogliflozin-induced vasodilation

The involvement of Kir channels in tofogliflozin-induced vasodilation was excluded using large-diameter femoral arteries, which do not express Kir channels^18, 19^. To assess the effect of other K^+^ channels in tofogliflozin-induced vasodilation, three distinct K^+^ channel inhibitors were applied to the femoral arterial rings. Pretreatment with the K_ATP_ channel inhibitor glibenclamide (10 μM) did not alter tofogliflozin-induced vasodilation (Figure 2A and B), nor did pretreatment with the BK_Ca_ channel inhibitor paxilline (1 μM) (Figure 2C and D). However, tofogliflozin-induced vasodilation was effectively inhibited in arterial rings pretreated with the Kv channel inhibitor 4-AP (3 mM) (Figure 2E and F). These results indicated that the vasodilatory effect of tofogliflozin is closely linked to the activation of Kv channels, but not to that of K_ATP_ or BK_Ca_ channels.

**Figure 2.**
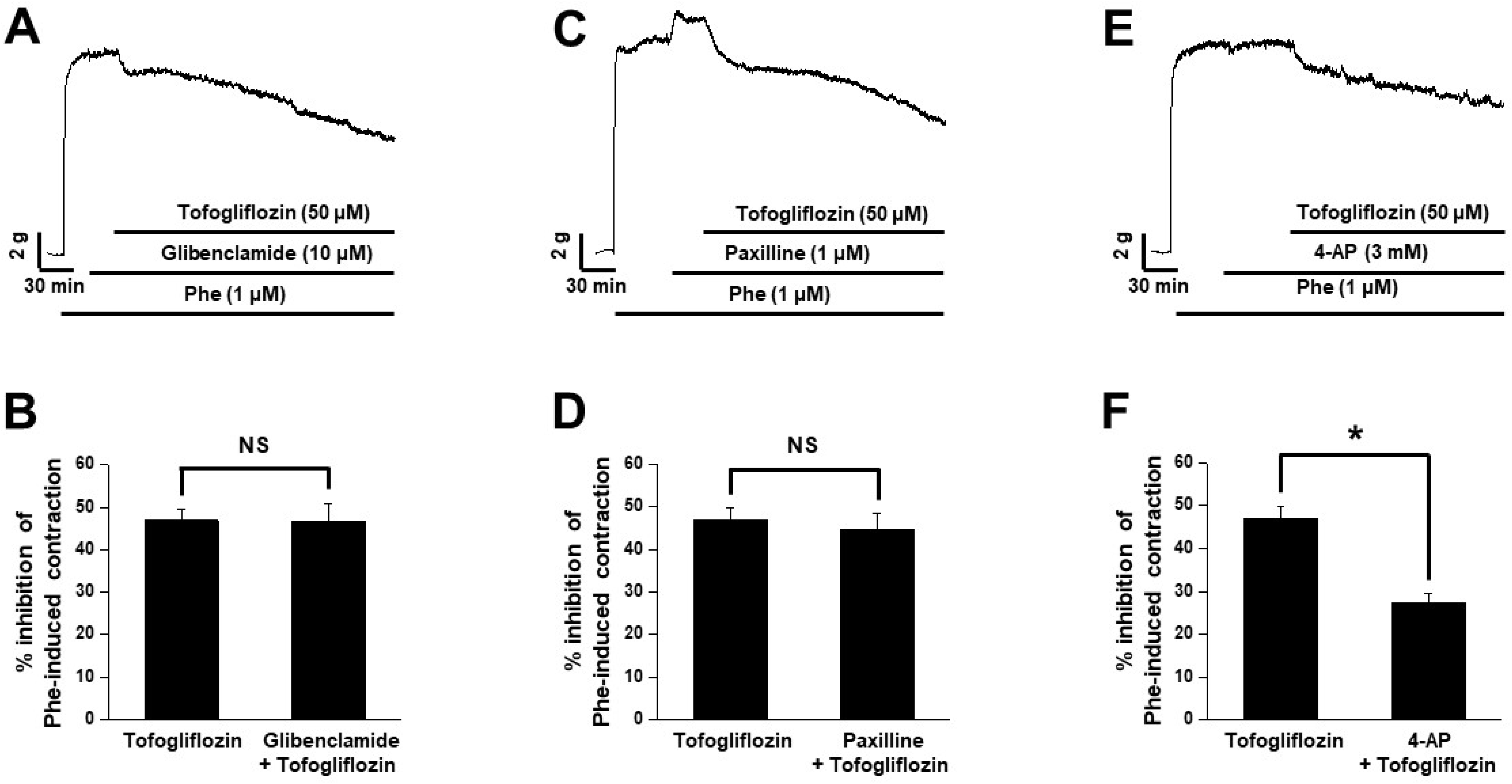
Involvement of K^+^ channels in tofogliflozin-induced vasodilation. (A) Vasodilatory effects of tofogliflozin in the presence of the K_ATP_ channel inhibitor glibenclamide. (B) Statistical analysis of the effect of glibenclamide on tofogliflozin-induced vasodilation (*n* = 8). NS = not significant. (C) Vasodilatory effects of tofogliflozin in the presence of the BK_Ca_ channel inhibitor paxilline. (D) Statistical analysis of the effect of paxilline on tofogliflozin-induced vasodilation (*n* = 5). NS = not significant. (E) Vasodilatory effects of tofogliflozin in the presence of the Kv channel inhibitor 4-AP. (F) Statistical analysis of the effect of 4-AP on tofogliflozin-induced vasodilation (*n* = 7). \**P* < 0.05.

### Effects of Kv1.5, Kv2.1, and Kv7.X inhibitors on tofogliflozin-induced vasodilation

The Kv subtype responsible for tofogliflozin-induced vasodilation was further investigated by pretreating the femoral arterial rings with inhibitors targeting the Kv1.5, Kv2.1, and Kv7.X channels prior to tofogliflozin administration. Pretreatment with the Kv1.5 channel inhibitor DPO-1 (1 μM) did not affect tofogliflozin-induced vasodilation (Figure 3A and B), nor did pretreatment with the Kv2.1 channel inhibitor stromatoxin-1 (100 nM) (Figure 3C and D). However, tofogliflozin-induced vasodilation was markedly reduced in arterial rings pretreated with the Kv7.X channel inhibitor linopirdine (10 μM) (Figure 3E and F). To confirm the involvement of the Kv7 subtype in the vasodilatory effect of tofogliflozin, Phe-induced pre-contracted femoral arterial rings were pretreated with another Kv7.X channel inhibitor, XE991 (10 μM). XE991 pretreatment also effectively reduced tofogliflozin-induced vasodilation (Figure 3G and H). These findings suggest that the activation of Kv channels, specifically the Kv7 subtype, plays a crucial role in tofogliflozin-induced vasodilation.

**Figure 3.**
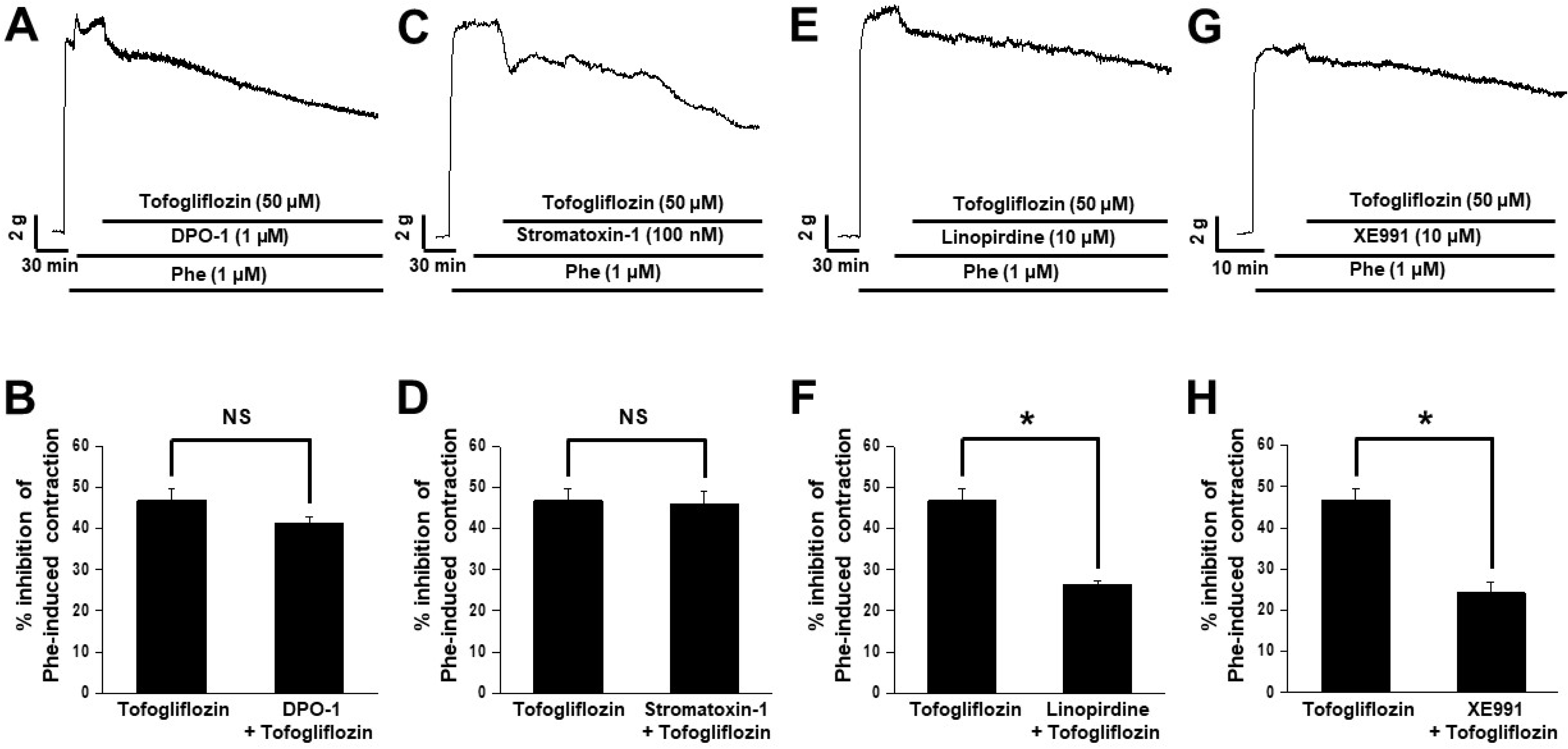
Involvement of Kv1.5, Kv2.1 and Kv7.X channel subtypes in tofogliflozin-induced vasodilation. (A) Vasodilatory effects of tofogliflozin in the presence of the Kv1.5 subtype inhibitor DPO-1. (B) Statistical analysis of the effect of DPO-1 on tofogliflozin-induced vasodilation (*n* = 5). NS = not significant. (C) Vasodilatory effects of tofogliflozin in the presence of the Kv2.1 subtype inhibitor stromatoxin-1. (D) Statistical analysis of the effect of stromatoxin-1 on tofogliflozin-induced vasodilation (*n* = 4). NS = not significant. Vasodilatory effects of tofogliflozin in the presence of the Kv7.X subtype inhibitors linopirdine (E) and XE991 (G). Statistical analysis of the effects of linopirdine (F) and XE991 (H) on tofogliflozin-induced vasodilation (*n* = 5). \**P* < 0.05.

### Effects of SERCA pump inhibitors on tofogliflozin-induced vasodilation

The close relationship between intracellular Ca^2+^ concentrations and vascular tension suggested a role for SERCA pumps in tofogliflozin-induced vasodilation. This was tested using two SERCA pump inhibitors: thapsigargin (1 μM) and cyclopiazonic acid (CPA) (10 μM). Pretreatment with thapsigargin significantly diminished the vasodilatory effect of tofogliflozin (Figure 4A and B). Pretreatment of the arterial rings with CPA yielded similar results (Figure 4C and D), which suggested that tofogliflozin-induced vasodilation is associated with SERCA pump activity.

**Figure 4.**
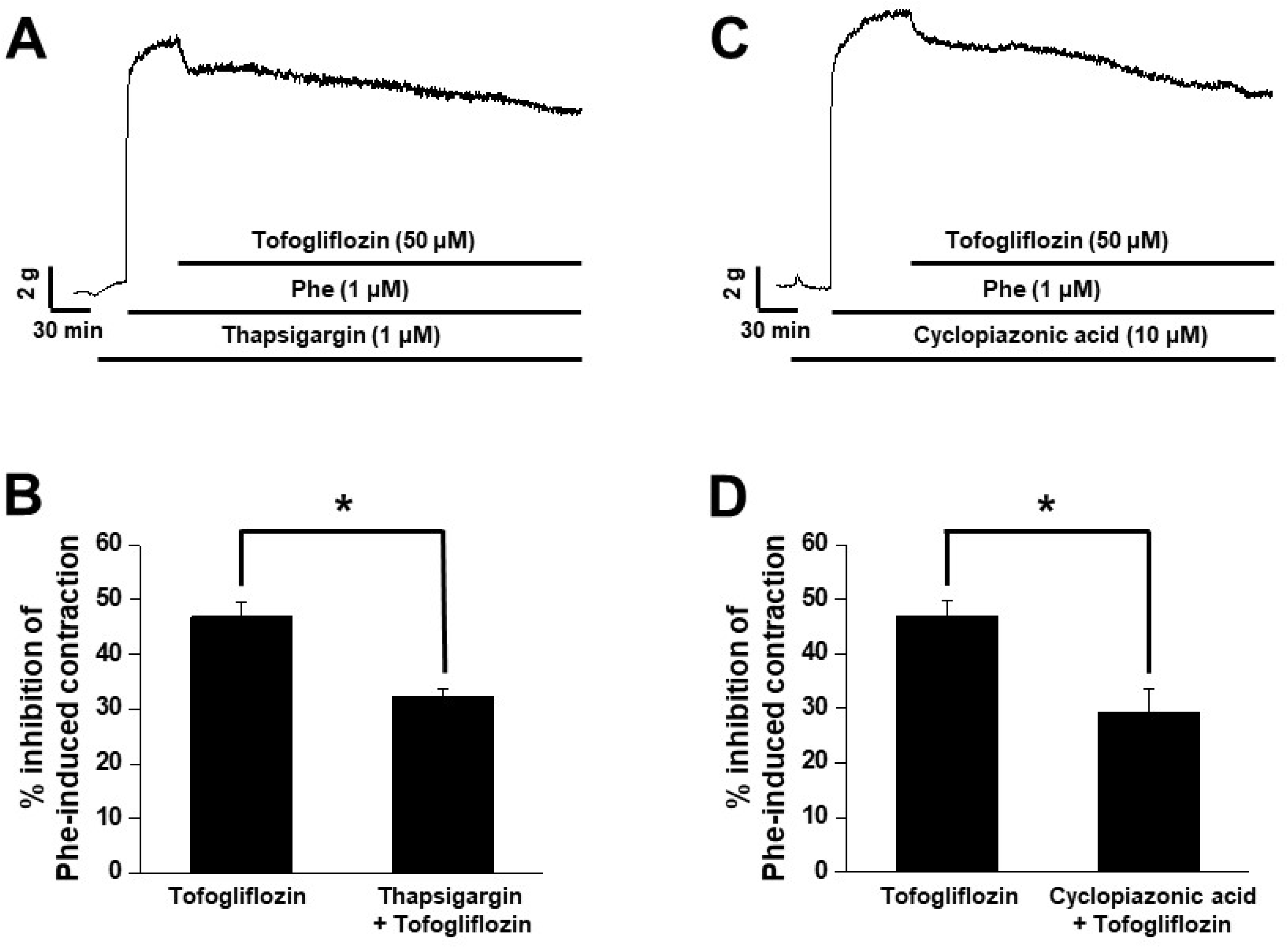
Involvement of sarco/endoplasmic reticulum Ca^2+^-ATPase (SERCA) pumps in tofogliflozin-induced vasodilation. (A) Vasodilatory effects of tofogliflozin in the presence of the SERCA pump inhibitor thapsigargin. (B) Statistical analysis of the effect of thapsigargin on tofogliflozin-induced vasodilation (*n* = 5). \**P* < 0.05. (C) Vasodilatory effects of tofogliflozin in the presence of another SERCA pump inhibitor cyclopiazonic acid (CPA). (D) Statistical analysis of the effect of CPA on tofogliflozin-induced vasodilation (*n* = 4). \**P* < 0.05.

### Effects of adenylyl cyclase (AC) and protein kinase A (PKA) inhibitors on tofogliflozin-induced vasodilation

Whether activation of the cAMP/PKA signaling pathways influences tofogliflozin-induced vasodilation was examined by treating the femoral arterial rings with the AC inhibitor SQ 22536 (50 μM) and the PKA inhibitor KT 5720 (1 μM). Pretreatment of the rings with SQ 22536 reduced intracellular cAMP production and thus PKA activity, but did not alter tofogliflozin-induced vasodilation (Figure 5A and B). Pretreatment with KT 5720 also had no effect (Figure 5C and D). These findings seemed to rule out the involvement of the cAMP/PKA signaling pathways in the vasodilatory response to tofogliflozin.

**Figure 5.**
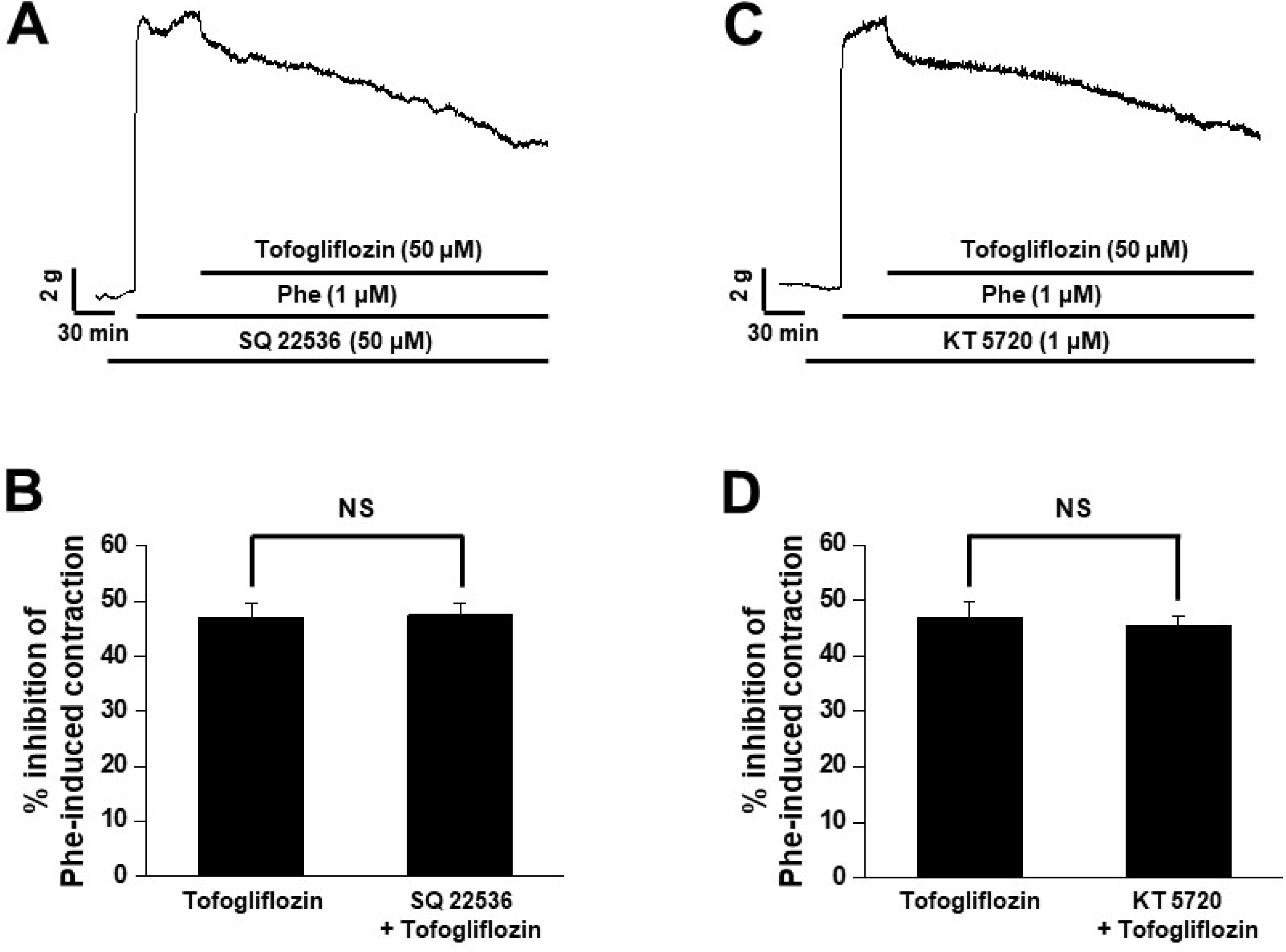
Absence of cAMP/protein kinase A (PKA) signaling pathway involvement in tofogliflozin-induced vasodilation. (A) Vasodilatory effects of tofogliflozin in the presence of the adenylyl cyclase (AC) inhibitor SQ 22536. (B) Statistical analysis of the effect of SQ 22536 on tofogliflozin-induced vasodilation (*n* = 5). NS = not significant. (C) Vasodilatory effects of tofogliflozin in the presence of the PKA inhibitor KT 5720. (D) Statistical analysis of the effect of KT 5720 on tofogliflozin-induced vasodilation (*n* = 5). NS = not significant.

### Effects of soluble guanylyl cyclase (sGC) and protein kinase G (PKG) inhibitors on tofogliflozin-induced vasodilation

Involvement of the cGMP/PKG signaling pathways in the vasodilation effect of tofogliflozin was determined using the sGC inhibitor ODQ (10 μM) and the PKG inhibitor KT 5823 (1 μM). Pretreatment of the arteries with ODQ prior to tofogliflozin administration significantly reduced tofogliflozin-induced vasodilation (Figure 6A and B), whereas KT 5823 pretreatment had no effect (Figure 6C and D). These findings indicated that tofogliflozin induces vasodilation by activating the sGC/cGMP signaling pathway, not the PKG pathway.

**Figure 6.**
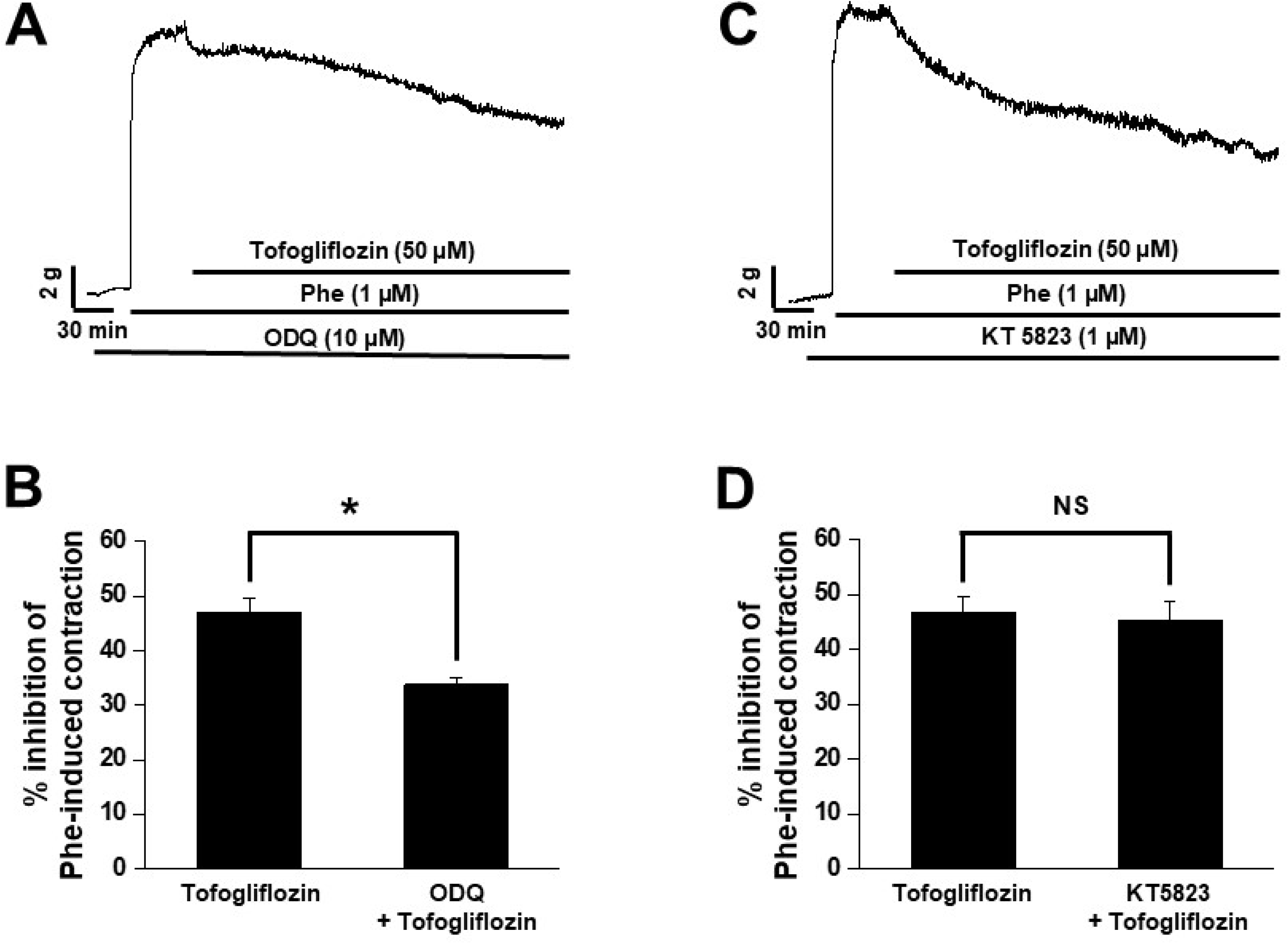
Involvement of the cGMP/ protein kinase A (PKG) signaling pathway in tofogliflozin-induced vasodilation. (A) Vasodilatory effects of tofogliflozin in the presence of the soluble guanyly cyclase (sGC) inhibitor ODQ. (B) Summary of the effects of ODQ on tofogliflozin-induced vasodilation. (*n* = 6). \**P* < 0.05. (C) Vasodilatory effects of tofogliflozin in the presence of the PKG inhibitor KT 5823. (D) Summary of the effects of KT 5823 on tofogliflozin-induced vasodilation. (*n* = 5). NS = not significant.

### Endothelium dependency of tofogliflozin-induced vasodilation

Given the crucial role of endothelial cell-derived factors in regulating vascular tension, the relationship between tofogliflozin-induced vasodilation and endothelial function was investigated. First, endothelial cells were removed from the femoral arterial rings by mechanically injecting bubbles, as described in the Materials and Methods. However, tofogliflozin-induced vasodilation was similar in femoral arterial rings with and without endothelium (Figure 7A and B). Application of the nitric oxide synthase (NOS) inhibitor L-NAME (100 μM) to endothelial-intact femoral arterial rings was used to assess the involvement of endothelial factors in tofogliflozin-induced vasodilation. Pretreatment with L-NAME did not alter the vasodilatory response to tofogliflozin (Figure 7C and D), nor did pretreatment with the low-conductance Ca^2+^-activated K^+^ channel (SK_Ca_) inhibitor apamin (1 μM) in combination with the intermediate-conductance Ca^2+^-activated K^+^ channel (IK_Ca_) inhibitor TRAM-34 (1 μM) (Figure 7E and F). These results demonstrated that the vasodilatory effects of tofogliflozin are independent of the endothelium.

**Figure 7.**
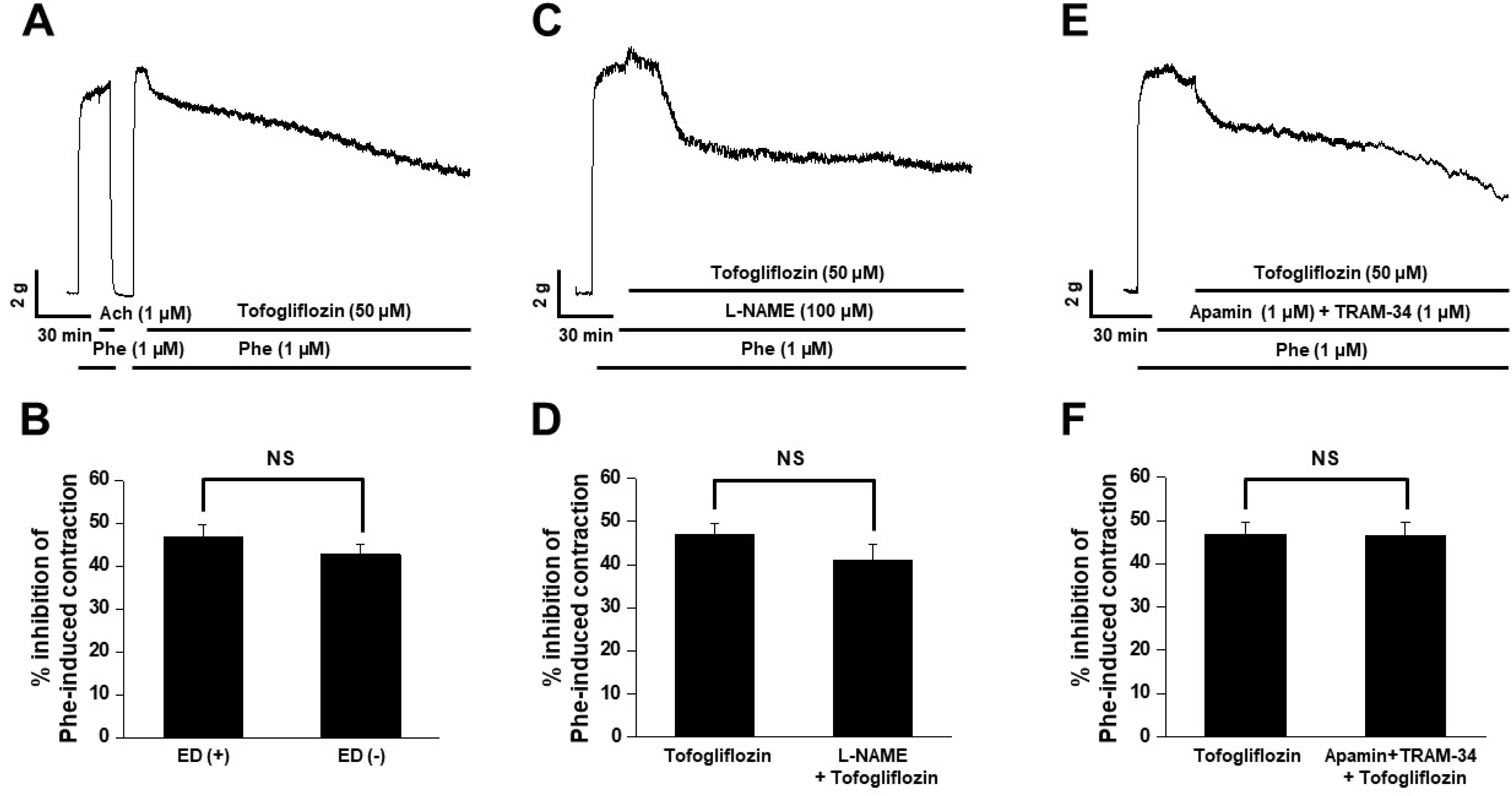
Involvement of the endothelium in tofogliflozin-induced vasodilation. (A) Vasodilatory effect of tofogliflozin on endothelium-denuded arteries. Successful removal of the endothelium was verified using acetylcholine. (B) Statistical analysis of tofogliflozin-induced vasodilation in endothelium-intact and -denuded arteries (*n* = 5). NS = not significant. (C) Vasodilatory effect of tofogliflozin in the presence of the nitric oxide synthase (NOS) inhibitor L-NAME. (D) Statistical analysis of the effect of L-NAME on tofogliflozin-induced vasodilation (*n* = 6). NS = not significant. (E) Vasodilatory effects of tofogliflozin in the presence of the SK_Ca_ inhibitor apamin and the IK_Ca_ inhibitor TRAM-34. (F) Statistical analysis of the effect of apamin + TRAM-34 on tofogliflozin-induced vasodilation (*n* = 5). NS = not significant.

### Effects of tofogliflozin on blood pressure

To assess the effects of tofogliflozin-induced vasodilation on blood pressure, changes in systolic and diastolic blood pressure were measured in rabbits administered tofogliflozin (1 mg/kg) via the ear vein. Both systolic and diastolic blood pressure decreased significantly in the tofogliflozin-treated rabbits (Figure 8A, B). Systolic blood pressure dropped from 126 to 85 mmHg, and diastolic blood pressure from 89 to 53 mmHg after tofogliflozin administration.

**Figure 8.**
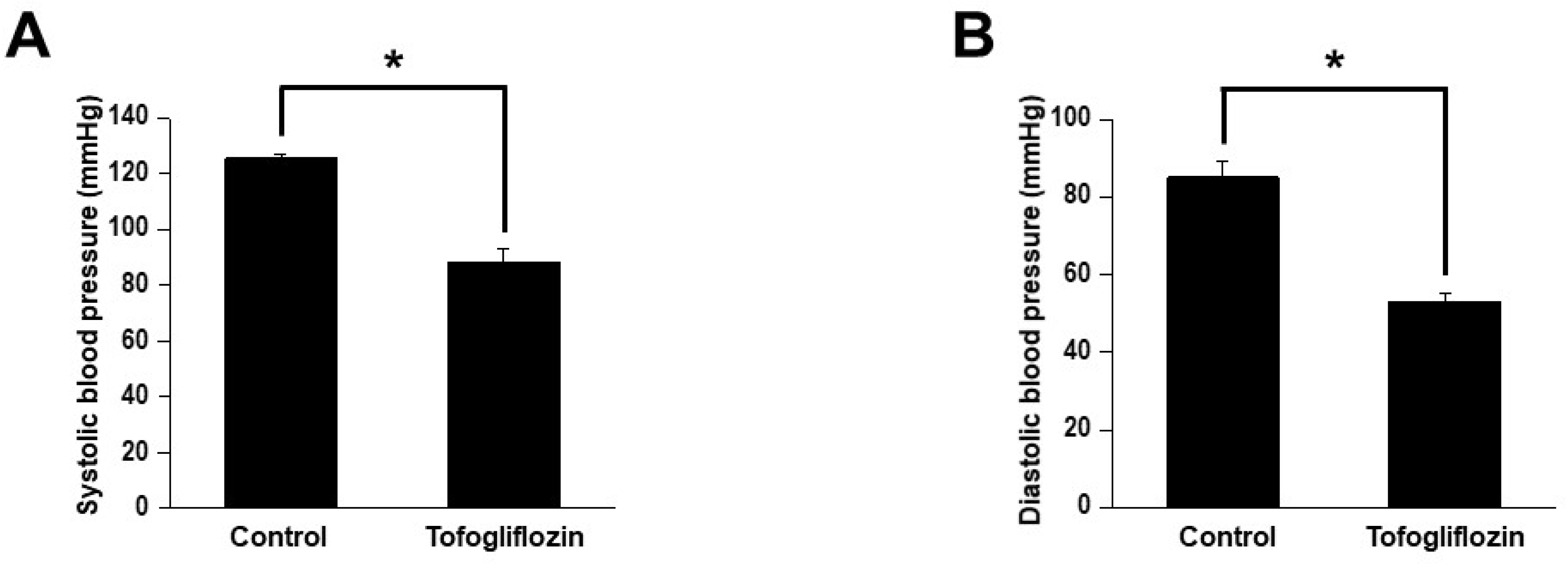
Effects of tofogliflozin on systolic and diastolic blood pressure. (A) Effects of tofogliflozin on systolic blood pressure (*n* = 4; *n* represents the number of rabbits). \**P* < 0.05. (B) Effects of tofogliflozin on diastolic blood pressure (*n* = 4; *n* represents the number of rabbits). \**P* < 0.05.

## DISCUSSION

This study showed that tofogliflozin induces concentration-dependent vasodilation in rabbit femoral arteries and that vasodilation is mediated by the activation of Kv channels, the cGMP signaling pathway, and the SERCA pump sensitization, rather than by the cAMP/PKA or PKG pathway, endothelium, or other K^+^ channels.

Nearly one third of T2DM patients worldwide are estimated to have cardiovascular disease, reflecting their 2–4-fold higher risk than the general population^20^. Therefore, the early selection and initiation of appropriate drug therapy is crucial to prevent the onset of cardiovascular disease in T2DM. SGLT2 inhibitors protect patients with T2DM from cardiovascular complications and are generally considered safe^21^. The sustained positive effects of the SGLT2 inhibitor tofogliflozin include reductions in hemoglobin A1c (HbA1c) and improvements in high-density lipoprotein cholesterol (HDL-C) levels, body mass index, and systolic blood pressure^22^. Clinical studies have reported significant reductions in the use of antihypertensive and lipid-lowering medications in patients treated with tofogliflozin compared with those treated with conventional therapy^23, 24^. However, the effect of tofogliflozin on vascular tension was not examined. Our study showed that tofogliflozin induces vasodilation (Figure 1) and identified the mechanisms underlying this effect.

K⁺ channels, including Kv, BK_Ca_, Kir, and K_ATP_, play a crucial role in the functional regulation of the vasculature^25–28^. Our study was able to link the vasodilatory effect of tofogliflozin to the activation of Kv channels, while ruling out roles for K_ATP_ and BK_Ca_ channels (Figure 2). Because Kir channels are not expressed in large-diameter arteries, they were excluded from our experiments. Kv channels exhibit significant inter- and intratissue heterogeneity, attributed to the diversity of their subtypes. Kv1.5, Kv2.1, and Kv7 are highly expressed in VSMCs and are involved in physiological functions such as maintaining resting membrane potential and regulating vascular tension, but also in pathological states. Kv1.5 channels in pulmonary VSMCs were shown to be upregulated by luteolin, a naturally occurring flavonoid, thereby promoting pulmonary vasodilation^29^; in diabetes, Kv2.1 channels are downregulated, resulting in increased arterial tone^30^. The medicinal plant rosemary activates Kv7 channels in VSMCs to induce vascular relaxation^31^. Thus, the complex roles of the Kv1.5, Kv2.1, and Kv7 subtypes in regulating vascular tone warrant further study. This study showed that tofogliflozin-induced vasodilation is mediated by activation of the Kv7 subtype, but not the Kv1.5 or Kv2.1 subtype (Figure 3). Further studies are needed to explore the broader implications of Kv7 activation by tofogliflozin, as well as the clinical applications of Kv7 as a therapeutic target in ensuring vascular health.

The SERCA pump is essential for maintaining intracellular Ca^2+^ levels and thus vascular tone^32^ as it transports intracellular Ca^2+^ into the sarco/endoplasmic reticulum, thereby reducing free Ca^2+^ concentrations and promoting vasodilation^33^. The role of the SERCA pump in regulating vasodilation has been investigated in several studies. For example, AMP-related kinase activation was shown to induce renal vasodilation via the SERCA pump in VSMCs^34^. Another study reported improved cardiac function and the prevention of cardiovascular diseases in association with increased SERCA pump activity^35^. The ability of both of the tested SERCA pump inhibitors, thapsigargin and CPA, to effectively inhibit the vasodilatory effect of tofogliflozin (Figure 4) suggests that tofogliflozin induces vasodilation by reducing intracellular free Ca^2+^ levels through SERCA pump activation. Therefore, the involvement of the SERCA pump should be considered in evaluations of the effects of particular drugs on the cardiovascular system.

Several therapeutic agents regulate vascular tone by acting on vascular K^+^ channels via the cAMP/PKA or cGMP/ PKG signaling pathways^36–38^. For example, braylin, a phosphodiesterase 4 inhibitor, induces potent vasodilation by increasing cAMP/cGMP levels, thereby activating BK_Ca_, Kir, and Kv channels^39^. In previous studies, we demonstrated an association between cAMP/PKA- or cGMP/PKG-related signaling pathways and vasodilation, as the SGLT2 inhibitors dapagliflozin and empagliflozin were shown to induce vasodilation by activating PKG and Kv channels^40, 41^. In the present study, we examined whether tofogliflozin-mediated vasodilation involved cAMP/PKA and/or cGMP/PKG-related signaling pathways by testing the effects of inhibitors of AC, sGC, PKA, and PKG. While the vasodilatory effect of tofogliflozin was not significantly inhibited in response to AC, PKA, and PKG inhibitors, it was significantly reduced by the sGC inhibitor ODQ (Figures 5 and 6). By promoting the generation of cGMP, sGC induces PKG activation; however, it can also influence cellular functions independently of PKG, by modulating ion channels and other proteins. For example, procyanidin C1-induced vasodilation is associated with the activation of the NO/cGMP pathway, which induces K^+^ channel activation^42^. Similarly, chrysin-induced vasodilation involves the NO/sGC/cGMP signaling cascade as well as K^+^ and Ca^2+^ channels^43^.

The regulation of vascular tone is not limited to smooth muscle factors but also involves endothelium-dependent dilation, NO synthase (NOS) activity, and NO signaling^44^. NOS is responsible for the production of NO, a well-established activator of sGC in VSMCs, such that NOS activation leads to increased cGMP production, ultimately resulting in vascular dilation^45^. Endothelial dysfunction may result in impaired endothelium-dependent vasodilation, a key factor in the development of diabetes-related cardiovascular diseases^46^. However, our results showed that the vasodilatory response of tofogliflozin did not differ between endothelium-denuded and endothelium-intact femoral arterial rings (Figure 7A, B). Consistent with these results, pretreatment with the NOS inhibitor L-NAME or the SK_Ca_ and IK_Ca_ inhibitors apamin and TRAM-34 did not affect tofogliflozin-induced vasodilation (Figure 7C–F). Our results demonstrate that the vasodilatory effects of tofogliflozin are not mediated by endothelium-dependent signaling pathways, but instead by interactions with Kv channels and the sGC/cGMP pathway in VSMCs. Further studies are needed to clarify the precise mechanisms.

Clinically, the recommended oral dose of tofogliflozin for adults is 20 mg daily^8^. Tofogliflozin has a short elimination half-life of 5–6 h and is primarily excreted in urine, which helps to minimize the risk of nocturnal hypoglycemia^22, 47^. In addition to its effect on glycemic control, the benefits of tofogliflozin include reductions in body weight and blood pressure, as well as improved HDL-C, triglyceride, and HbA1c levels^22, 48^. The standard tofogliflozin concentration of 50 μM was used in this study. This was higher than the maximum plasma concentration (1.32 μM)^47^, as *in vitro* experiments require a higher dose than *in vivo* measurements, but lower tofogliflozin concentrations, ranging from 1 to 30 μM, have also been shown to induce vasodilation. Given the high input resistance of vascular smooth muscle, even small changes in vascular tension can lead to significant changes in blood pressure or flow. Although tofogliflozin shows potential in treating both hypertension and diabetes, its vasodilatory effects are an important clinical concern in patients with low blood pressure.

## CONCLUSION

In summary, this study showed that tofogliflozin induces vasodilation by activating SERCA pumps, sGC/cGMP pathways, and Kv channels, leading to a reduction in blood pressure. There was no evidence supporting the involvement of the endothelium, cAMP/PKA, PKG-related pathways, or other K^+^ channels in the effect of tofogliflozin. Our elucidation of the mechanisms responsible for the vascular protective effects of tofogliflozin will allow its therapeutic potential to be maximized and unintended risks to be avoided.

## Abbreviations

T2DM: Type 2 diabetes mellitus
SGLT2: Sodium-glucose cotransporter 2
NOS: Nitric oxide synthase
SK_Ca_: Small-conductance Ca²⁺-activated K⁺ channels
IK_Ca_: Intermediate-conductance Ca²⁺-activated K⁺ channels
Kv: Voltage-dependent K^+^
K_ATP_: ATP-sensitive K^+^
BK_Ca_: Large-conductance Ca2+-activated K^+^
SERCA: Sarcoplasmic/endoplasmic reticulum Ca^2+^-ATPase
CPA: Cyclopiazonic acid
PKA: Protein kinase A
PKG: Protein kinase G
sGC: soluble guanylyl cyclase
VSMCs: Vascular smooth muscle cells
cAMP: cyclic adenosine monophosphate
cGMP: cyclic guanosine monophosphate

## Acknowledgments

This work was supported by the National Research Foundation of Korea (NRF) grant funded by the Korea government (RS-2023-00239172). This work was also supported by the China Scholarship Council (202209307001).

